# Phenotypic and genomic analysis of *P* elements in natural populations of *Drosophila melanogaster*

**DOI:** 10.1101/047910

**Authors:** I.A. Kozeretska, V. Bondarenko, V.I. Shulga, S.V. Serga, A.I. Rozhok, A.V. Protsenko, M.G. Nelson, C.M. Bergman

## Abstract

The *Drosophila melanogaster P* transposable element provides one of the best cases of horizontal transfer of a mobile DNA sequence in eukaryotes. Invasion of natural populations by the *P* element has led to a syndrome of phenotypes known as “P-M hybrid dysgenesis” that emerges when strains differing in their *P* element composition mate and produce offspring. Despite extensive research on many aspects of *P* element biology, questions remain about the stability and genomic basis of variation in P-M dysgenesis phenotypes. Here we report the P-M status for a number of populations sampled recently from Ukraine that appear to be undergoing a shift in their *P* element composition. Gondal dysgenesis assays reveal that Ukrainian populations of *D. melanogaster* are currently dominated by the P’ cytotype, a cytotype that was previously thought to be rare in nature, suggesting that a new active form of the *P* element has recently spread in this region. We also compared gondal dysgenesis phenotypes and genomic *P* element predictions for isofemale strains obtained from three worldwide populations of *D. melanogaster* in order to guide further work on the molecular basis of differences in cytotype status across populations. We find that the number of euchromatic *P* elements per strain can vary significantly across populations but that total *P* element numbers are not strongly correlated with the degree of gondal dysgenesis. Our work shows that rapid changes in cytotype status can occur in natural populations of *D. melanogaster*, and informs future efforts to decode the genomic basis of geographic and temporal differences in *P* element induced phenotypes.

## Introduction

A substantial portion of eukaryotic genomes is represented by transposable elements (TEs). These TE families include those that colonized genomes long ago during the evolution of the host species and groups, but also those that have appeared in their host genomes recently. The genome of *Drosophila melanogaster* is thought to have been colonized by the *P* element family of transposons at least 70 years ago as a result of a horizontal transmission event from *D. willistoni* [1, 2], a species that inhabits South America, the Caribbean and southern parts of North America. Laboratory strains of *D. melanogaster* established from wild populations before the 1950s did not contain *P* element, while by the late 1970s this TE family was found in all populations worldwide [1]. The specific molecular and evolutionary mechanisms that promoted the global spread of the *P* element over such a short period are not still clearly understood.

The presence of *P* elements induces a number of phenotypes in *Drosophila* that can be characterized by the so-called “P-M hybrid dysgenesis” assay [3]. Among the most prominent *P* element induced phenotypes is gonadal dysgenesis (GD), which is the key marker determining P-M status in particular strains of flies [1, 4–10]. In the P-M system, fly strains can be classified by their phenotypes as follows: P-strains have both *P* element transposition inducing and repressing abilities, P’-strains only have the inducing ability, Q-strains only have the repressing ability, and M-strains have neither [3–4, 11]. An M-strain carrying some *P* sequences in the genome is called M’ to distinguish it from true M-strains, which are completely devoid of *P* elements [12]. Hybrid dysgenesis phenotypes were previously thought to be controlled by repressor proteins of various kinds that are encoded by *P* elements of different sizes [13]. More recent work has shown that these phenotypes mostly arise due to RNA interference mechanisms mediated by piRNAs produced by telomeric *P* elements, such as TP5 and TP6, and the effects are amplified by RNAs produced by other *P* elements [14].

Natural populations of *D. melanogaster* are currently thought to have a relatively stable distributions of *P* elements and P-M status. For example, Australian populations demonstrate a north-south cline of the frequency of various *P* element copies [6], which underwent only minor changes in the frequencies of truncated and full-size copies of the *P* element a decade later [13]. In North American populations, P and Q cytotypes prevailed in the 1970s-1980s [1, 5]. Two decades later, no significant changes were detected, except for an increase in the proportion of truncated *P* element copies [15]. A similar phenotypic stability has also been observed in populations from Eurasia, Australia, Japan and Taiwan [7-8, 16]. Cytotype status has also remained stable for over 30 to 40 years in some Japanese populations [9, 17–18]. Some authors propose that these protracted periods of stability follow the initial transition from M cytotype to P, Q and M’ cytotypes [9].

Studies of *P* element insertions at the molecular level provide the best opportunity to characterize the dynamics of this TE in natural populations. Such analyses have been hampered by the lack of unbiased and practical methods to characterize the frequencies of *P* element insertions at individual sites across large numbers of strains. In the past, estimates of TE insertion frequencies were obtained by *in situ* hybridization to polytene chromosomes [19–23]. However, the resolution of *in situ* hybridization is limited, with highly truncated insertions being missed entirely and nearby insertions not being distinguished from one another. More recent studies have used targeted PCR to survey populations for TE insertions identified in the reference genome sequence [24–28]. These surveys are necessarily biased towards insertions with high population frequencies [24], and are not applicable in the case of the *P* element since the ISO-1 strain used for the reference genome sequence is an M strain lacking any *P* elements [28].

Application of whole-genome next generation sequencing (NGS) technologies has improved many areas of population genetics research. To date, hundreds of re-sequenced genomes of *D. melanogaster* exist and can be freely used for population and genomic analyses [29–33]. Moreover, a number of computational algorithms have been designed for *de novo* TE insertion discovery, annotation, and population analysis in *Drosophila* [34–40]. Comparison of different methods for detecting TEs in *Drosophila* NGS data has shown that they identify different subsets of TE insertions [41]. Determining which TE detection method is best for specific biological applications remains an area of active research.

Here we report the recent P-M status of a number of populations from Ukraine sampled in 2012-2013. Previous studies of flies collected in Ukraine from 1936 through the 1980’s revealed only M and M’ cytotypes [1, 42–43]. More recent collections in Ukraine from 2008 and 2009 revealed only M’ cytotypes [44], consistent with the stability in cytotype status observed in other regions around the world. Here we present evidence that the P’ cytotype is now prevalent in several locations in Ukraine, suggesting a relatively rapid recent shift in the cytotype status of these populations. To better understand the molecular basis for differences in cytotype status among populations, we also compared GD phenotypes and genomic *P* element predictions for isofemale strains obtained from three worldwide populations of *D. melanogaster.* This analysis revealed that the number of euchromatic *P* elements per strain can vary significantly across populations but is not correlated with the degree of GD. Finally, we show that ability to predict *P* elements in pooled population samples is dependent on the number of strains pooled, indicating that efforts to rigorously detect differences in the number of *P* elements across populations using pool-seq must control for read depth per strain. Our work shows that rapid changes in cytotype status can occur in natural populations of *D. melanogaster,* and informs future efforts to decode the genomic basis of differences in *P* element induced phenotypes over time and space.

## Materials and Methods

Sample collection.Flies were collected at fruit processing facilities and in apple orchards in late August to early September in 2012 and 2013 at seven locations in Ukraine: Uman’ (48°45'45.26"N – 30°14'38.97"E), Varva (50°29'33.30"N – 32°42'50.93"E), Inkerman (44°61'77.64"N – 33°61'44.14"E), Chornobyl (51°16'13.73"N – 30°13'19.63"E), Kyiv (50°21'9.06"N – 30°28'57.70"E), Poliske (51°14'9.49"N – 29°23'51.60"E), Chornobyl Nuclear Power Plant (NPP) (51°22'30.01"N – 30°8'21.33"E). Single gravid females were used to establish isofemale strains. For each population, isofemale strains collected from one year were used for analysis.

### Gonadal dysgenesis phenotyping

The cytotype of a particular isofemale strain was determined by the GD assay, for which two types of crosses were used: cross A (M-strain (Canton S) females x tested males) and A* (tested females x P-strain (Harwich) males) [4, 42]. Three A and three A* crosses were routinely made for each isofemale strain and 50 F1 females of each cross were dissected for gonadal status inspection. Both unilateral and bilateral ovary reduction was counted. The GD score for each isofemale strain was calculated using the formula %GD =½(%GD1) + %GD2, where %GD1 stands for the percent of individuals with unilateral reduction of the ovary, and %GD2 means the percent of individuals with bilateral gonadal reduction. P-M characteristics were defined as in [5]. The result of GD tests for each population was shown as a two-dimensional graph, plotting A %GD versus A* %GD, following [45].

In addition to collecting GD phenotypes for populations from Ukraine, we also re-analyzed GD assay data from [46]. These cytotype data are for isofemale strains from three geographic regions: North America (Athens, Georgia, USA), Europe (Montpellier, France), and Africa (Accra, Ghana), described in [47]. Definitions of A and A* crosses in [46] are inverted relative to those proposed by [4], and were standardized prior to re-analysis here.

### Genome-wide identification of P element insertions

Whole genome shotgun sequences described in [31] from the three populations analyzed by [46] [North America (Athens, Georgia, USA), Europe (Montpellier, France), and Africa (Accra, Ghana)] were downloaded from the European Nucleotide Archive (ERP009059). These data were collected using a uniform library preparation and sequencing strategy (thus mitigating many possible technical artifacts) and include data for both individual isofemale strains and pools of isofemale strains [31]. Two pool-seq samples were analyzed for each population [North America (15 and 30 strains), Europe (20 and 39 strains), and Africa (15 and 32 strains)].

*P* element insertions were identified by TEMP (revision d2500b904e2020d6a1075347b398525ede5feae1; [38]) and RetroSeq (revision 700d4f76a3b996686652866f2b81fefc6f0241e0; [48]) using the McClintock pipeline (revision 3ef173049360d99aaf7d13233f9d663691c73935; [49]). McClintock was run across the major chromosome arms (chr2L, chr2R, chr3L, chr3R, chr4, chrY, and chrX) of the dm6 version of the Release 6 reference genome [50] using the following options: ‐C ‐m "retroseq temp" ‐i ‐p 12 ‐b. Reference TE insertions needed for TEMP were generated automatically by McClintock using RepeatMasker (version open-4.0.6). The *D. melanogaster* TE library used by McClintock to predict reference and non-reference TE insertions is a slightly modified version of the Berkeley *Drosophila* Genome Project TE data set v9.4.1 described in [51, 52].

In addition to providing "raw" output for each component method in the standardized zero-based BED6 format, McClintock generates "filtered" output tailored for each method. TEMP output was filtered to (i) eliminate predictions where the start or end coordinates had negative values, (ii) retain predictions where there is sequence evidence supporting both ends of an insertion, and (iii) retain predictions that have a ratio of reads supporting the insertion to non-supporting reads of >10%. Likewise, RetroSeq predictions were filtered to (i) eliminate predictions where two different TE families shared the same coordinates and (ii) retain predictions assigned a call status of greater than or equal to six as defined by [53].

Graphical and statistical analyses were performed in the R programming environment (version 3.2.1).

## Results

### Cytotype status of natural populations from Ukraine

To determine cytotypes in natural populations of *D. melanogaster*, we carried out GD assays for 241 isofemale strains from 7 locations in Ukraine (Table 1, Fig. 1, File S1). We identified two main cytotypes in these populations, namely M [12] and P’ [11]; strains with Q or P cytotype were not detected. Previous work suggests M-strains detected here are actually likely to be M’ [44]. Interestingly, the majority of lines were of P’ (86%; 207/241), a cytotype which has not been previously reported in this region and is thought to be rare in nature [1, 42–44]. Most populations demonstrated a higher proportion of P’-strains relative to M’-strains, except for the population from Inkerman, which was equally represented with P’-strains and M’-strains.

Differences in GD phenotypes for isofemale strains among Ukrainian populations were observed in both inducing ability (One-way ANOVA; F=44.5, 6 d.f., P<2e-16) and susceptibility (One-way ANOVA; F=160.46, 6 d.f., P<2e-16) (Fig. S1). The average inducing ability of a strain varied from 11.0% (Inkerman) to 51.1% (Poliske) in A crosses of the GD test. However, in most populations (Uman’, Kyiv, Chornobyl, Varva), the inducing ability values were much closer to each other (21.4-26.4%). The greatest range in inducing ability within a population was observed from Chornobyl, in which the %GD varied from 4-65%.

**Table 1.** Gondaldysgenesis levels and P-M status for isofemale strains of *D. melanogaster* obtained from natural populations in Ukraine. Previous work suggests M-strains reported here are likely to be M’ [44].

| Population | Year of collection | Cross A (%GD±SD) | Cross A* (%GD±SD) | # strains | M | Q | P' | P |
| --- | --- | --- | --- | --- | --- | --- | --- | --- |
| Uman' | 2012 | 21.4±6.9 | 83.1±8.5 | 38 | 3 | 0 | 35 | 0 |
| Kyiv | 2012 | 24.9±6.4 | 84.4±7.5 | 49 | 2 | 0 | 47 | 0 |
| Chornobyl | 2012 | 24.7±15.0 | 52.5±16.3 | 40 | 5 | 0 | 35 | 0 |
| Inkerman | 2013 | 11.0±4.9 | 24.4±9.2 | 23 | 11 | 0 | 12 | 0 |
| Poliske | 2013 | 51.1±8.6 | 73.7±13.2 | 18 | 0 | 0 | 18 | 0 |
| Chornobyl NPP | 2013 | 13.6±6.6 | 75.2±8.2 | 34 | 12 | 0 | 22 | 0 |
| Varva | 2013 | 26.4±8.8 | 35.8±11.0 | 39 | 1 | 0 | 38 | 0 |
| Total |  |  |  | 241 | 34 | 0 | 207 | 0 |

**Figure 1.**
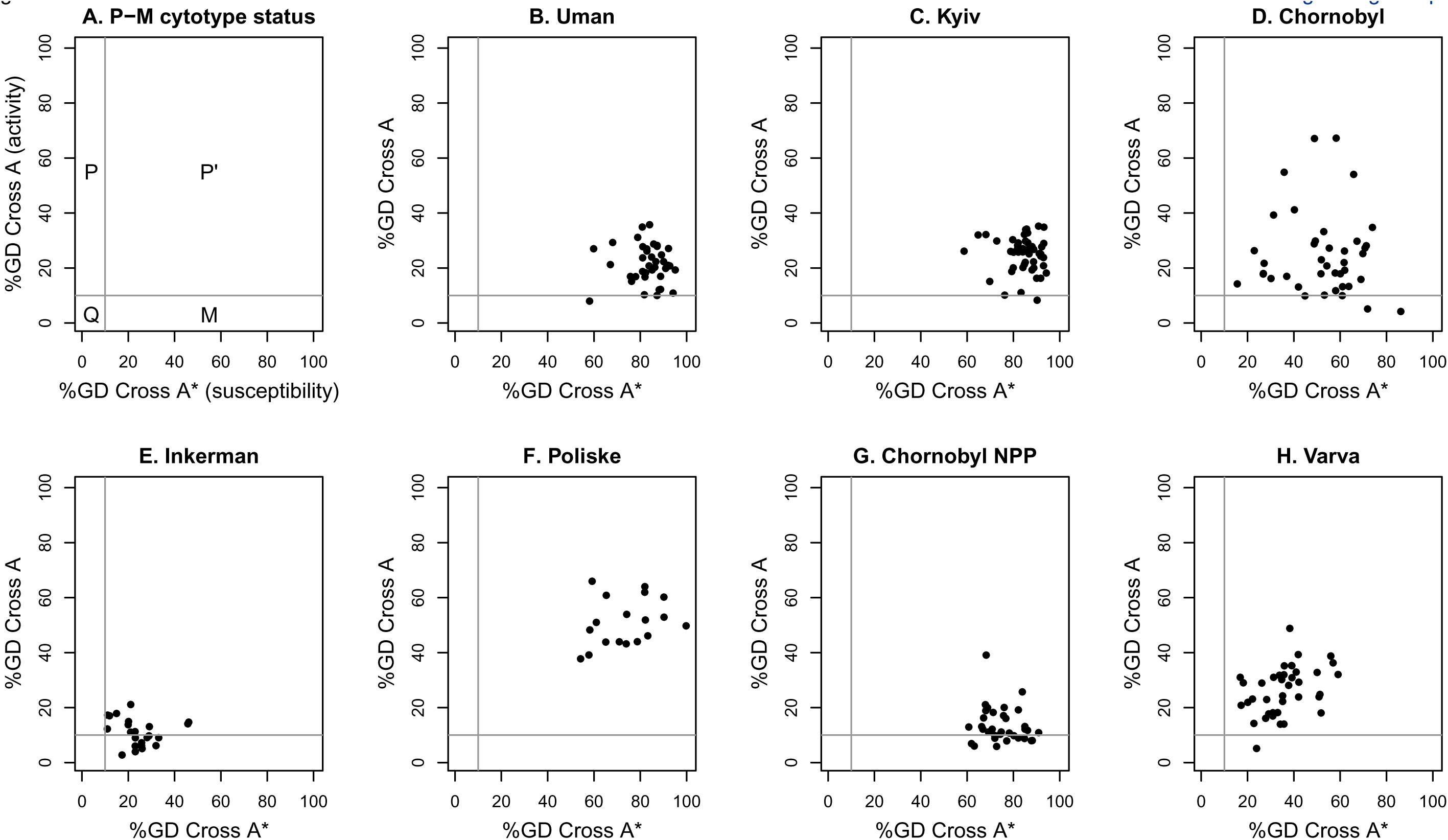
Results of GD assays for isofemale strains from natural populations from Ukraine.The graphs present %GD for the populations for cross A (vertical axis) and cross A* (horizontal axis). Each dot represents an isofemale strain. The P-M status for various sectors of GD phenotypic space defined by A and A* crosses are according to [44] and shown in panel A.

The average susceptibility to *P* transposition varied from 24.4% (Inkerman) to 84.5% (Kyiv) in A* crosses of the GD test. In most populations (Uman’, Kyiv, Chornobyl NPP, Poliske), the susceptibility values were very high (>73.7%), with Chornobyl and Varva having a more intermediate average susceptibility. We did not detect strains with susceptibility below 10%. For A* crosses, the greatest range in susceptibility was observed in the population from Chornobyl.

Overall, most populations from Ukraine demonstrated a moderate ability to induce *P* transposition, with moderate to high susceptibilities to *P* transposition. One exception to this rule is for the population from Inkerman, which demonstrated low inducing abilities and low susceptibility. In general, all the populations showed less variation in inducing ability (A crosses) relative to susceptibility (A* crosses). In addition, most strains from each population clustered together reasonably well in phenotypic space, with the exception of the Chornobyl population, which exhibited substantial variation for both GD phenotypes.

### Comparison of cytotype status and P element insertions in individual strains from North America, Europe and Africa

Answering questions about how cytotype status varies geographically and temporally requires a better understanding of how genomic *P* element insertion profiles relate to *P* element induced phenotypes. As a preliminary step towards this goal, we identified *P* element insertions in publicly available genomic data [31] for a panel of 43 isofemale strains from three global regions with previously-published GD phenotypes [46]. In contrast to the Ukrainian populations reported here, isofemale strains from these populations were mainly P-strains, M-strains and Q-strains (Fig. 2, Table 2, File S2). According to our genomic analysis below, strains that are defined phenotypically M are actually M’ strains. Additionally, strains from these populations do not differ significantly in the degree of their inducing ability (One-way ANOVA; *F*=0.06, 2 d.f., *P*=0.94) or susceptibility (One-way ANOVA; *F*=1.66, 2 d.f., *P*=0.2) (Fig. S2A–B).

**Figure 2.**
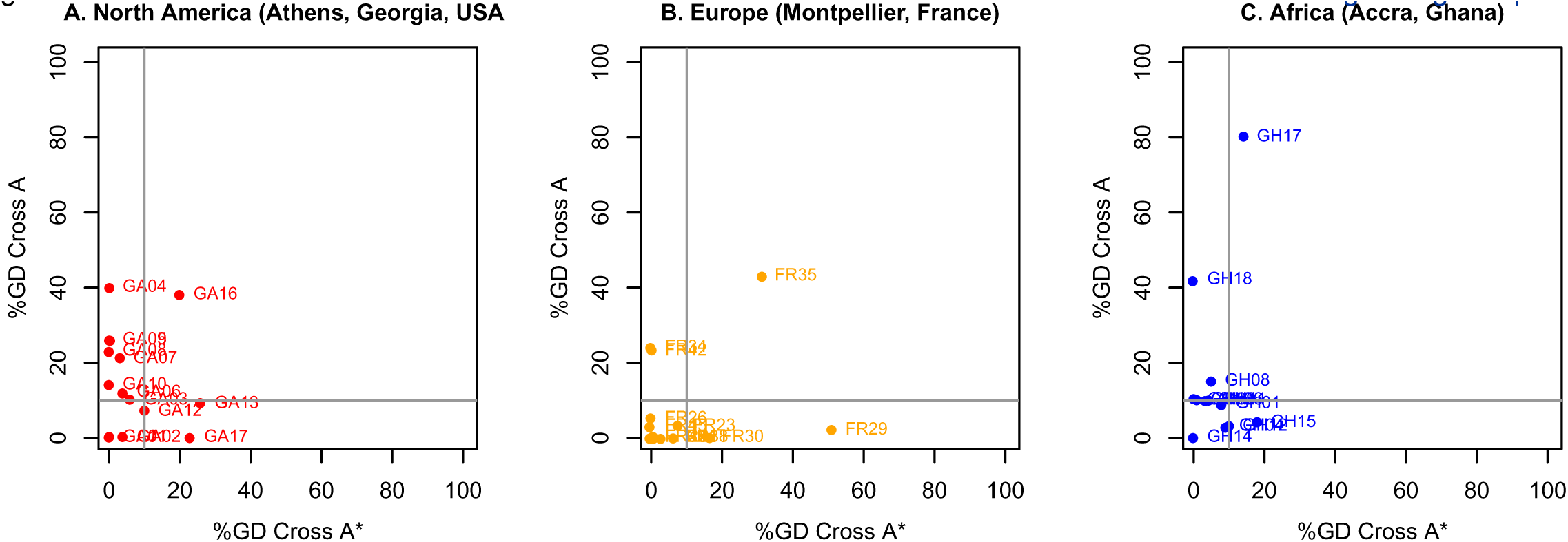
Results of GD tests for isofemale strains from natural populations from North America, Europe and Africa.The graphs present %GD for the populations for cross A (vertical axis) and cross A* (horizontal axis) using data from [45]. A and A* cross definitions in [45] are inverted relative to those proposed by [4] and were standardized prior to analysis here. Each dot represents an isofemale strain. Definition of P-M status by A and A* crosses according to [44] is the same as in Fig. 1A.

**Table 2.**
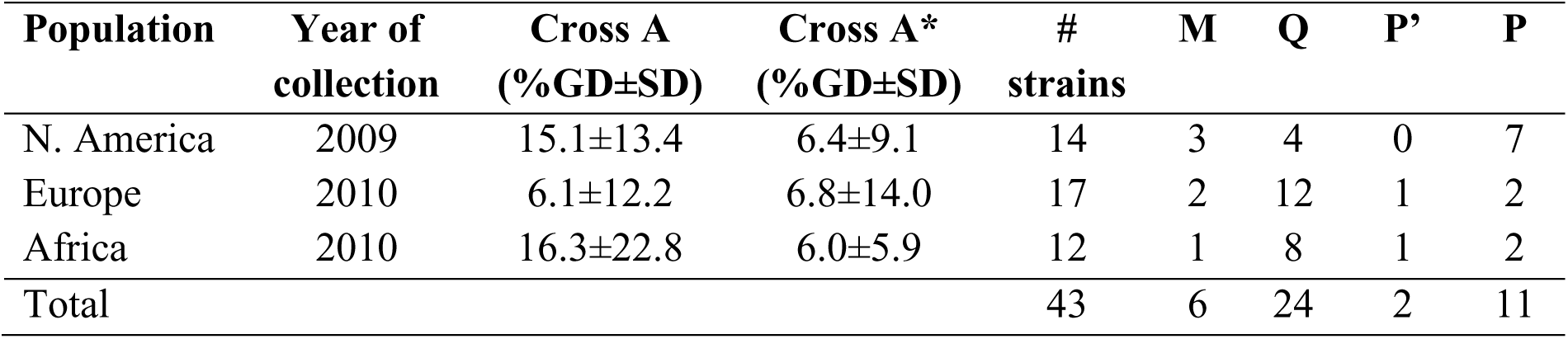
Gondaldysgenesis levels and P-M status for isofemale strains of *D. melanogaster* obtained from natural populations in North America, Europe and Africa. Phenotypically M-strains are in fact M’-strains based on analysis of genomic data (File S2, File S3).

We predicted *P* element insertions in the genomes of these strains using two independent bioinformatics methods – TEMP and RetroSeq – to ensure that our conclusions are not dependent on the idiosyncrasies of a single TE detection software package (File S2, File S3). We also investigated the effects of the standard filtering of TEMP and RetroSeq output performed by McClintock, a meta-pipeline that runs and parses multiple TE insertion detection methods. Across all three populations, the numbers of *P* elements predicted per strain were well correlated across strains, regardless of the method of analysis and filtering (*r*≥0.712) (Fig. S3). The correlation in the number of *P* elements predicted per strain was highest for the filtered TEMP and filtered RetroSeq datasets (*r*=0.945), suggesting that the post-processing steps performed by McClintock improve the consistency of TE predictions made by these methods on isofemale strains. For filtered datasets, the number of *P* elements predicted per strain was 52–134 for TEMP and 52–162 for RetroSeq.

We tested whether the number of *P* elements per strain is associated with GD phenotypes and found neither cross A nor A* to be significantly linearly correlated with the filtered number of predictions made by TEMP or RetroSeq (p>0.11; Fig. 3). Similar results were obtained for the raw output of these methods as well (Fig. S4). However, we did find evidence that strains from the North American populations carry fewer euchromatic *P* element insertions in their genomes relative to the European and African populations, regardless of the TE detection method and filtering (One-way ANOVA; F>9.26; 2 d.f., P<5e-4) (Fig. 3; Fig. S2C–F). These results suggest that there is no simple relationship between the number of euchromatic*P* elements and GD phenotypes at the level of individual strains, and that systematic differences in the abundance of *P* elements per strain may not lead to differences in the frequency of P-M phenotypes at the population level.

**Figure 3.**
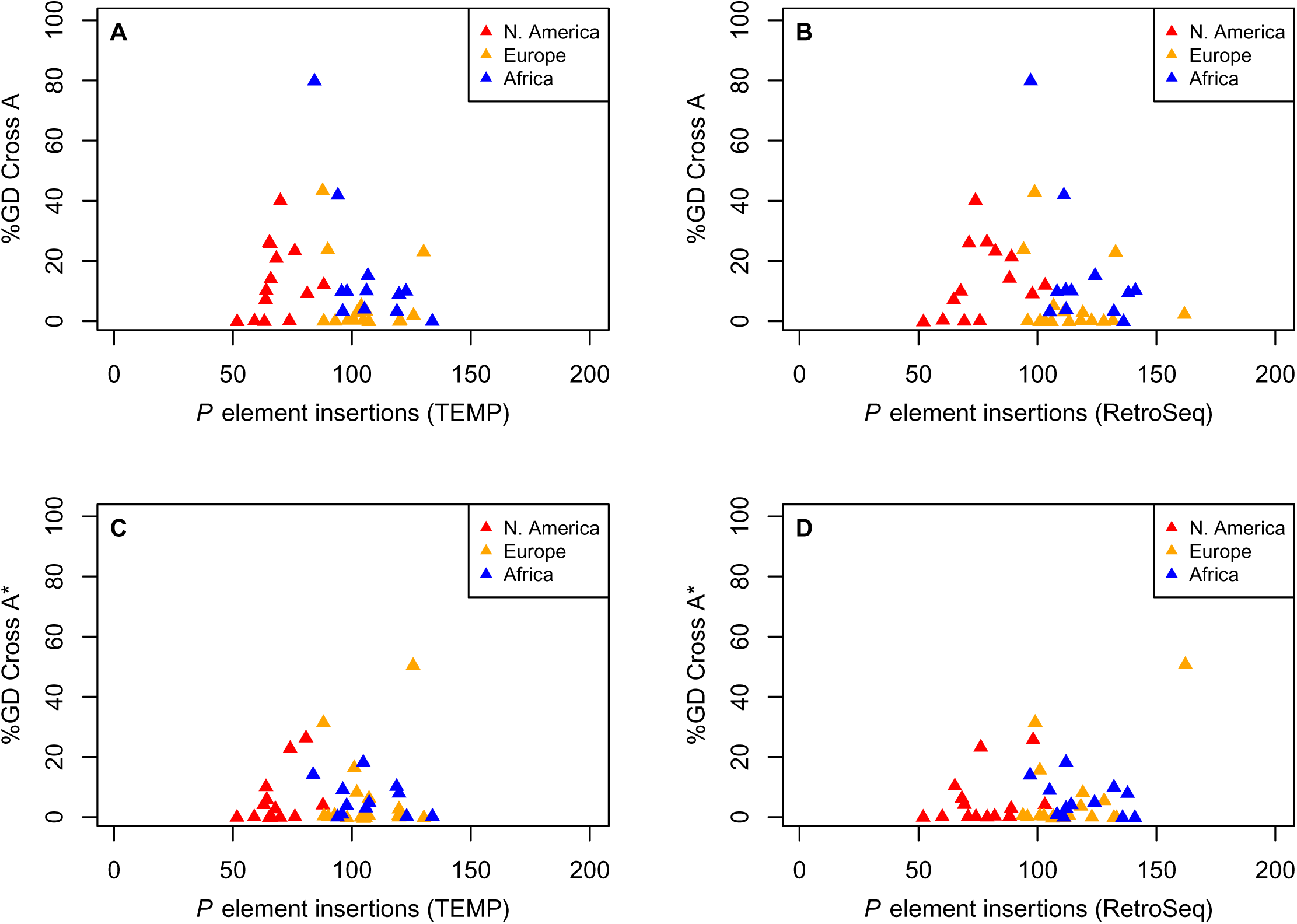
Relationship between %GD in A and A* crosses and numbers of euchromatic*P* element insertions identified by TEMP or RetroSeq for isofemale strains from natural populations from North America, Europe and Africa. %GD data are from [45] and are the same standardized values as in Figure 2. Numbers of *P* elements predicted by TEMP or RetroSeq shown here are after standard filtering by McClintock (see Materials and Methods for details). Results for unfiltered raw output of TEMP or RetroSeq are shown in Fig. S4. Each triangle represents an isofemale strain.

### Population difference in P element insertion numbers can be observed in pool-seq samples

Pooled-strain sequencing (pool-seq) is a cost-effective strategy to sample genomic variation across large numbers of strains and populations [54]. To address whether the differences among populations we observed in the number of *P* elements predicted in isofemale strain data are also seen in pool-seq data, we predicted *P* element insertions in pool-seq samples from the same populations (Table 3, File S3). Two pool-seq samples are available for each population that differ in the number of individuals/isofemale strains used: North America (n=15 and n=30), Europe (n=20 and n=39), and Africa (n=15 and n=32). The smaller pools from each population include one individual from the same isofemale strains analyzed above; the larger pools contain one individual from the same strains as the smaller pools, plus individuals from additional isofemale strains from the same population that were not sequenced as isofemale strains. Thus, the smaller pool samples are a nested subset of the larger pool samples, and pool-seq samples from the same population are not fully independent from one another. However, larger and smaller strain pool samples contain similar numbers of reads (~44 million read pairs per sample).

**Table 3.**
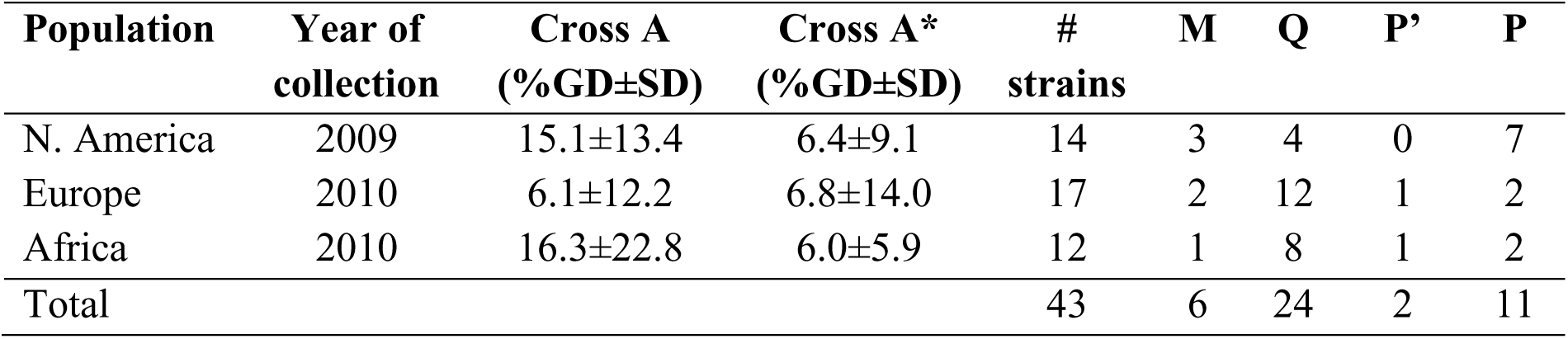
Numbers of *P* element insertions identified by TEMP and RetroSeq in pool-seq samples from three worldwide populations of *D. melanogaster*. Columns labeled raw and filtered represent output generated by each method before or after standard filtering by McClintock, respectively (see Materials and Methods for details).

The numbers of *P* element insertions identified by TEMP and RetroSeq in these pool-seq samples are given in the Table 3. For the raw output, TEMP predicted more insertions in the larger strain pools relative to the smaller strain pools, as expected for a method designed to capture TE insertions that are polymorphic within a sample [38]. However, the McClintock-filtered TEMP output included many fewer insertions per strain pool than in the raw output, as well as fewer insertions in the larger strain pools relative to the smaller strain pools. These effects are likely because of the requirement for default McClintock filtering of TEMP predictions to have a ratio of reads supporting the insertion to non-supporting reads of >10%. McClintock filtering reduced the total number of RetroSeq predictions only by less than 2-fold, with fewer insertions being predicted for the larger strain pools in both raw and filtered RetroSeq output. Regardless of method or filtering, when the number of strains in a pool-seq sample was used as a scaling factor, fewer insertions were predicted per strain in the larger strain pools relative to smaller strain pools for all of the populations studied. Because smaller pools contain a subset of the same strains that are present in larger pools, these results suggest a dilution effect whereby, at a fixed sequencing coverage, *P* element insertions that are predicted in smaller pools cannot be detected in larger pools, even though they are in fact present in the sample.

In spite of this dilution effect, African pool-seq samples tended to have more insertions per strain than North American samples as seen in the isofemale strain datasets. This result is most clearly demonstrated for the comparison between North American and African samples with 15 strains where African sample has more predicted insertions regardless of TE detection method and filtering. These results suggest that, despite dilution effects, pool-seq samples can capture general population trends in total *P* element insertion numbers seen in isofemale strain sequencing.

## Discussion

We analyzed the P-M status of strains from seven natural populations of *D. melanogaster* from Ukraine. Strains from one of these populations (Uman’) have a long history of study (starting from 1966) [1]. Another population from Magarach (Crimea) was represented with isofemale strains collected from the wild in 1936 [1, 42–43]. Rozhok et al (2009) most recently characterized the PM status of nine populations from Ukraine and revealed only the M’ cytotype, which is consistent with data from Europe [7]. Thus, flies collected in Ukraine in from the 1930s to 2009 appeared to have harbored only M or M’ cytotypes. The results presented here indicate that Ukrainian populations of *D. melanogaster* are currently dominated by the P’ cytotype, a cytotype that was previously thought to be rare in nature but has recently been found in the southern USA and Turkey [10, 15]. The sudden prevalence of the P’ cytotype in these populations suggests that a new active form of the *P* element has recently spread in Ukraine.

Several studies before 2000 describe P-M status as phenotypically stable in populations from Japan, Eurasia, Africa, Oceania, and North America [1, 4–10]. The apparent rapid changes in the P-M status in Ukraine do not follow this this general pattern, however the mechanisms underlying these changes remain unclear. It has been suggested that high activity of TEs may result from the stresses that accompany adaptation to new climates [55]. Such a response to stress has been suggested for populations with M’ cytotype [15] which has previously been found in Ukrainian populations [44].

We also performed an exploratory analysis of genomic data from three worldwide populations of *D. melanogaster* with published GD phenotypes [31, 46–47]. The main aim of this analysis was to inform future computational and sampling strategies to decode the genomic basis of variation in *P* element induced phenotypes such as those observed in Ukraine. However, our results allowed us to draw several preliminary conclusions about the detection of *P* element insertions in *D. melanogaster* population genomic data and the genomic basis of GD phenotypes.

For samples derived from isofemale strains, we find that two different TE detection methods (TEMP and RetroSeq) generate well-correlated numbers of *P* element predictions per strain, but that filtering by McClintock improves the overall correlation between these methods (mainly by reducing the number of presumably false positive TEMP predictions). Analysis of pool-seq samples revealed larger differences between TE detection methods and a larger effect of McClintock filtering. Regardless of method or filtering, we found that there is a diminishing return on the number of *P* element insertions detected per individual/strain in pool-seq samples for a given sequencing coverage. This dilution effect means it is difficult to compare numbers of predicted *P* element across pool-seq samples unless the number of strains and/or the read depth is carefully controlled. Diminishing returns per strain also implies there are allele frequency threshold effects for calling *P* element insertions in NGS samples, which may also impact attempts to integrate results from pool-seq data with those based on isofemale strains.

Regardless of prediction method, we found no simple linear correlation between the strength of GD phenotypes and the number of euchromatic *P* element insertions across isofemale strains (Fig. 3). Strains with very different number of euchromatic *P* elements appear to be able to generate very similar phenotypic outcomes. These results may imply that GD phenotypes are determined by specific full-length *P* element insertions that are active rather than overall numbers that include many inactive defective copies, or that GD phenotypes are controlled by heterochromatic *P* element insertions not detected here. Alternatively, the lack of correlation between the number of *P* element insertions and GD phenotypes may imply that the genomic sequence data is not of sufficient depth in these samples, or that the bioinformatic methods are not sensitive enough to detect all *P* element insertions in an isofemale strain and introduce noise that obscures a true relationship.

We did however observe differences in the number of predicted *P* element insertions at the population level (Fig. 3), even though we found no strong differences in the levels of GD phenotypes across these populations. Specifically, the North American population had the fewest predicted *P* element insertions, regardless of the TE detection method or filtering (Fig. 3, Fig. S2). Evidence for fewer *P* element insertions per strain in the North American population could also be detected in pool-seq samples, especially when the number of strains is controlled for (Table 3). Thus it is possible to detect clear differences in euchromatic*P* element insertion profiles among populations using either isofemale strains or pool-seq, however interpreting how these profiles relate to P-M phenotypes at the strain or population level remains an open challenge.

In conclusion, based on the increase in a novel P’ cytotope, we suggest that at the present time there is a new *P* element type that is active in Ukrainian populations of *D. melanogaster*. Further analysis of genomic data from these populations is needed to understand the dynamics of this invasion and connect variation in GD phenotypes to *P* element insertions at the molecular level.

## Acknowledgements

This work was supported by Wellcome Trust PhD Studentship (096602/B/11/Z) to MGN and a Human Frontier Science Program Young Investigator grant RGY0093/2012 to CMB.

Authors declare no competing interests.

## Supplemental Files

**File S1.**
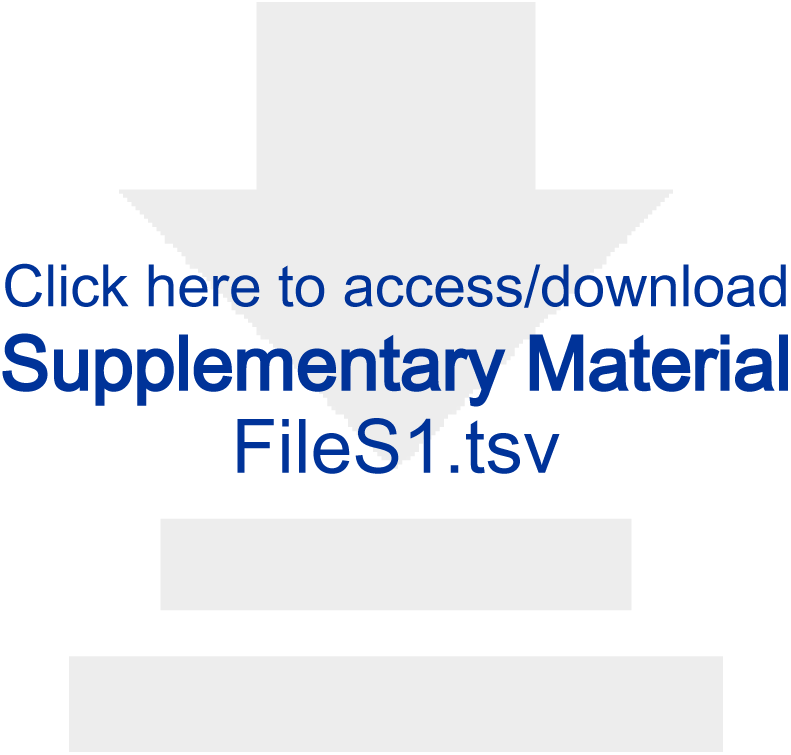
Tab separated value formatted file with %GD data from A and A* crosses, P–M cytotype status, and population for isofemale strains from Ukraine.

**File S2.**
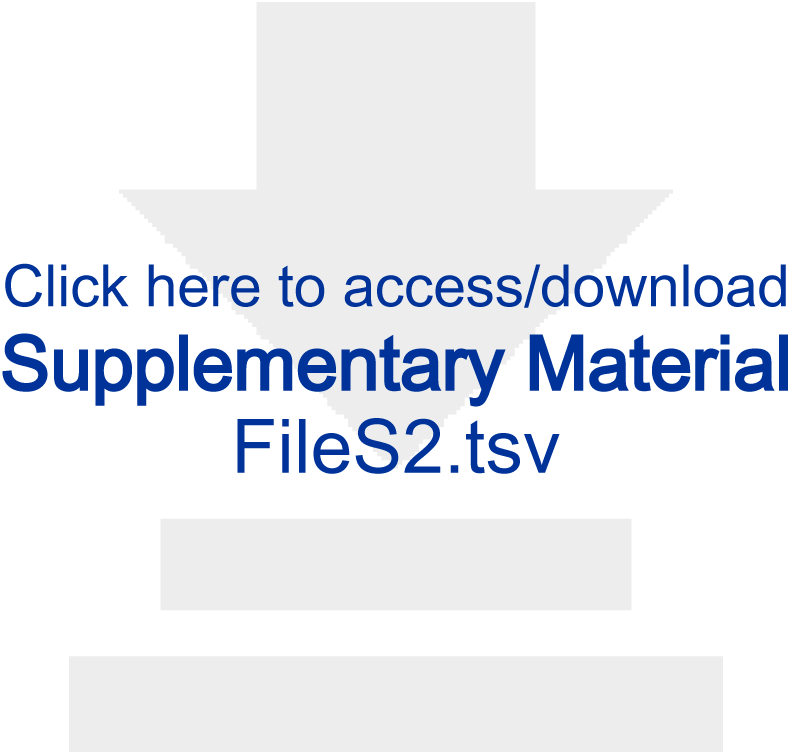
Tab separated value formatted file with %GD data from A and A* crosses, P–M cytotype status, population, and numbers of predicted *P* elements in raw and filtered output from TEMP and RetroSeq, respectively, for isofemale strains from three global regions. GD data are taken from [46] and were standardized to definitions proposed by [4] prior to re-analysis here.

**File S3.**
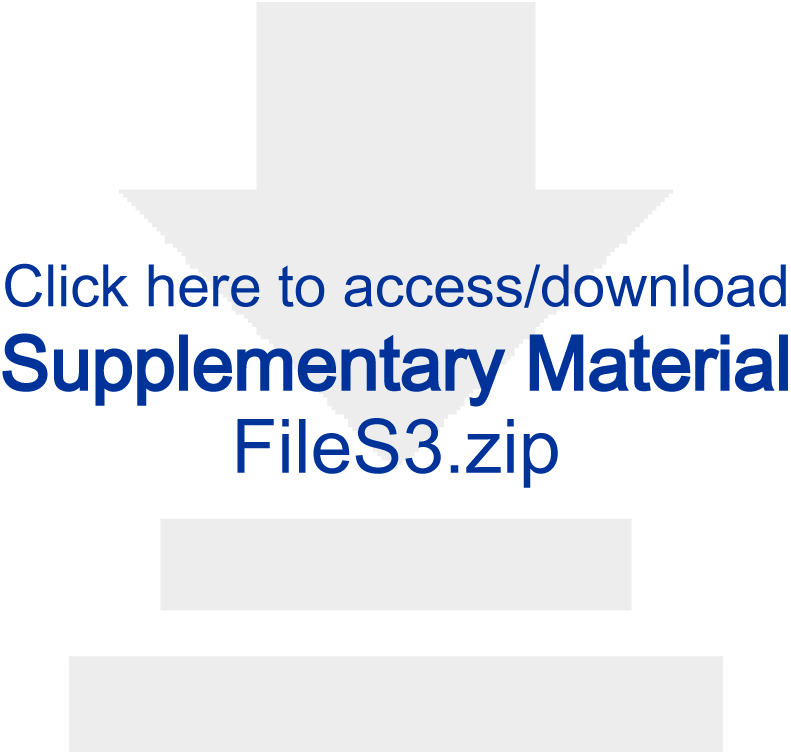
Zip archive of browser extensible data (BED) files of predicted *P* element locations in genome sequences from 50 isofemale strains and 6 pool-seq samples from three global regions. Each sample has four BED files corresponding to raw (*raw.bed) and filtered (*nonredundant.bed) output from TEMP and RetroSeq, respectively.

**Figure S4.** Relationship between %GD in A and A* crosses and numbers of euchromatic *P* element insertions identified by TEMP or RetroSeq for isofemale strains from natural populations from North America, Europe and Africa. %GD data are from [45] and are the same standardized values as in Figure 2. Numbers of *P* elements predicted by TEMP or RetroSeq shown are raw output prior to standard filtering by McClintock. Results for McClintock-filtered output of TEMP and RetroSeq are shown in Fig. 3. Each circle represents an isofemale strain.

